# phiC31 integrase for recombination mediated single copy insertion and genome manipulation in *C. elegans*

**DOI:** 10.1101/2020.11.25.398784

**Authors:** Fang-Jung Yang, Chiao-Nung Chen, Tiffany Chang, Ting-Wei Cheng, Ni-Chen Chang, Chia-Yi Kao, Chih-Chi Lee, Yu-Ching Huang, Shih-Peng Chan, John Wang

**Affiliations:** Biodiversity Research Center, Academia Sinica, Taipei, Taiwan; Genome and Systems Biology Degree Program, College of Life Science, National Taiwan University, Taipei 10617, Taiwan; Graduate Institute of Microbiology, College of Medicine, National Taiwan University, Taipei 10051, Taiwan

## Abstract

*C. elegans* benefits from a large set of tools for genome manipulation. Yet, the insertion of large DNA constructs and the generation of inversions is still challenging. Here, we adapted the phiC31 integrase system for *C. elegans.* We generated an integrated phiC31 integrase expressing strain flanked by attP sites that also serves as a landing pad for integration of transgenes by recombination mediated cassette exchange (RCME). This strain is *unc-119(-)* so RMCE integrants can be produced simply by injection of a plasmid carrying attB sites flanking *unc-119(+)* and the gene(s) of interest. Additionally, phiC31 integrase is removed concomitantly with integration, eliminating the need to outcross away the integrase. Integrations are relatively easy to obtain for insert sizes up to ~15 kb. Taking advantage of this integration method we establish a dual color fluorescent operon reporter system to study post-transcriptional regulation of mRNA. Last we show that large chromosomal segments can be inverted using phiC31 integrase. Thus the phiC31 integrase system should be a useful addition to the *C. elegans* toolkit.

## Introduction

Genomic manipulations are a cornerstone of modern experimental biology. Molecular genetic analysis relies on the ability to create precise mutations, transgenes, and genomic rearrangements such as inversions or translocations. *C. elegans* benefits from a large set of genetic tools to do this (Stinchcomb *et al.* 1985; Mello *et al.* 1991; Way *et al.* 1991; Praitis *et al.* 2001; Frokjaer-jensen *et al.* 2008; Frokjaer-jensen *et al.* 2014; Dickinson and goldstein 2016; Iwata *et al.* 2016; Dejima *et al.* 2018; Nance and frokjaer-jensen 2019; Nonet 2020).

In some cases, precise single copy insertions, possibly of large size, are necessary or desired in *C. elegans.* Currently, this can be achieved through MosSCI (Frokjaer-jensen *et al.* 2008), CRISPR/Cas9 followed by repair (Dickinson and goldstein 2016), and Flp/Frt (Nonet 2020). Single or low copy insertions can also be created using biolistic bombardment (Praitis *et al.* 2001) and miniMos (Frokjaer-jensen *et al.* 2014), but the insertion sites for both methods cannot be controlled. A limitation of the MosSCI and CRISPR/Cas9 methods is that inserts longer than ~2 kb are difficult to obtain (Dickinson and goldstein 2016). The Flp/FRT system can insert transgenes of sizes in the 10-20 kb range relatively easily through recombination mediated cassette exchange (RMCE). While insert sizes greater than 20 kb may be possible, this has not been tested. Additionally, because Flp recombinase recombines two identical FRT sites, the products are also FRT sites. Consequently, an inserted a sequence may be excised.

phiC31 integrase is a phage-derived site-specific recombinase that has been used for genomic manipulations in *E. coli* (Merrick *et al.* 2018), yeast (Xu and brown 2016), flies (Groth *et al.* 2004; Bateman *et al.* 2006; Venken *et al.* 2006), mouse (Belteki *et al.* 2003), humans (Groth *et al.* 2000), fish (Hu *et al.* 2011; Lu *et al.* 2011; Kirchmaier *et al.* 2013), and plants (Thomson *et al.* 2010; Bernabe-orts *et al.* 2020; Cody *et al.* 2020). phiC31 integrase recombines an attB and an attP site to create an attR and an attL site. Recombination is a unidirectional reaction, so excision of integrated DNA is not a concern. phiC31 integrase has been used for “simple” insertion of donor constructs, for example by recombination of a single attB site on a plasmid with a single attP landing site in the genome, and for RCME between a pair of oppositely oriented attB sites on a plasmid with a pair of oppositely oriented attP sites in the genome. In addition, phiC31 can be used to delete DNA segments that are flanked by an attB and an attP site in the same orientation or invert DNA segments if the two sites are in opposite orientation. Despite its potential utility, limited tests to establish a phiC31 integrase system have been unsuccessful or partially successful in *C. elegans* (Vargas 2012; Frokjaer-jensen *et al.* 2014).

We had several motivations for developing the phiC31 integrase system in *C. elegans.* First, we were having difficulties inserting moderately large sequences (~5-15kb) with the available techniques. Larger constructs also allow for the conducting of more complex transgenic experiments, such as multigene constructs, especially those with longer genes or longer regulatory regions. Another reason was to be able to create precise very large insertions (>10 kb). This capability would allow the better characterization of a case of non-Mendelian inheritance of chromosomes in *C. elegans* (and *Caenorhabditis* species), whereby fathers that are heterozygous for chromosomes with a length difference, transmit longer chromosomes preferentially to sons (Wang *et al.* 2010; Le *et al.* 2017). Additionally, an efficient phiC31 integrase system could allow the testing of multiple transgene variants at the same genomic context faster. A final motivation is synthetic biology. phiC31 and other site-specific recombinases have been used to invert or delete DNA segments as both components and outputs in synthetic recorders and logic circuits (Merrick *et al.* 2018). The vast majority of synthetic biology experiments have been done in single cellular organisms, but could be applied to animals.

In this study we adapted the phiC31 integrase system for *C. elegans.* We first used MosSCI to insert a RMCE “landing pad” strain that also expresses phiC31 integrase and demonstrated that we could generate precise RMCE insertions. Then, we improved the landing pad strain such that RMCE integrants can be screened by phenotypic rescue of Unc into wild-type (WT) worms. Using this improved strain, we show that the insertion of large constructs is efficient. Taking advantage of this we establish a dual color fluorescent operon reporter system to study post-transcriptional regulation of mRNA. Last we show that large chromosomal segments can be inverted using phiC31 integrase. Thus the phiC31 integrase system should be a useful addition to the *C. elegans* toolkit.

## Materials and Methods

### Strains

N2,

BRC0189 *(ttTi5605 II; unc-119(ed9) III),*

BRC0546 *(antIs30 [attP-f; Cb-unc-119(+); glh-2p::phiC31; rol-6(partial); myo2p::gfp; attP-r] II; unc-119(ed9) III*),

BRC0566 *(antIs31 [attP-f; Cb-unc-119(ant40); glh-2p::phiC31; rol-6(partial); myo2p::gfp; attP-r] II; unc-119(ed9) III),*

BRC0587 *(dpy-13(e184) unc-30(ok613)),*

BRC0609 *(antSi50* [attB in *dpy-13] IV),*

BRC0664 (*antIs31 II*),

BRC0699 (*antSi51* [attP in *unc-30*] *IV*),

BRC0671 *(antSi50 antSi51),*

BRC0682 (*antIn1* [*dpy-13(ant41) unc-30(ant42)*]),

SPN369 *(chsIs001 [attR-f; SCMp::GFP(PEST)::H2B::lin-41 3′UTR::SL2::mCherry::H2B::unc-54 3′UTR; unc-119(+)] II; unc-119(ed9) III*),

SPN377 *(chsIs002 [attR-f; SCMp::GFP(PEST)::H2B::lin-41 3′UTRΔLCS::SL2::mCherry::H2B::unc-54 3′UTR; unc-119(+)] II; unc-119(ed9) III),*

SPN391 *(chsIs001 [attR-f; SCMp::GFP(PEST)::H2B::lin-41 3′UTR::SL2::mCherry::H2B::unc-54 3′UTR; unc-119(+)] II; unc-119(ed9) III; let-7(n2853) X*),

SPN393 *(chsIs002 [attR-f; SCMp::GFP(PEST)::H2B::lin-41 3′UTRΔLCS::SL2::mCherry::H2B::unc-54 3′UTR; unc-119(+)] II; unc-119(ed9) III let-7(n2853) X*),

SPN401 *(chsIs004 [attR-f; dpy-30p::GFP(PEST)::H2B::lin-41 3′UTR::SL2::mCherry::H2B::unc-54 3′UTR; unc-119(+)] II; unc-119(ed9) III)*

### Generation of the phiC31 integrase expressing landing pad strain

A plasmid containing the landing pad construct was cloned using the 4 fragments in the MultiSite Gateway Pro Plus system (ThermoFisher, cat no. 12537100). The first donor fragment contained a 221 bp attP site and *Caenorhabditis briggsae unc-119(+).* The second fragment contained the germline *glh-2* promoter, phiC31 integrase, and the *glh-2* 5-UTR. phiC31 was codon optimized and has three artificial introns and an SV40 NLS on the C-terminus. The third fragment is composed of a partial *rol-6* gene. Originally the dominant *rol-6(su1006)* allele was intended both as a marker for MosSCI integration and as a negative selection marker for successful RMCE. However, we failed to obtain such a Rol strain through MosSCI, so a truncated donor fragment was made instead to be compatible with the 4-fragment Gateway cloning. The fourth fragment contained *myo-2p::gfp* (enhanced GFP, but hereafter GFP) and an attP site. The attP sites are arranged in “opposite” orientation in the landing pad construct.

MosSCI (Frokjaer-jensen *et al.* 2008) was used to integrate the phiC31 integrase construct at the *ttTi5605* site on LG II. After screening >10,000 P0 equivalents, we obtained one integrant (BRC0546(*antIs30*)), which we verified by a combination of PCR assays and sequencing of amplification products. Expression and splicing of phiC31 was also confirmed by RT-PCR.

To obtain an Unc-119 strain, the *Cb-unc-119(+)* gene within the *antIs30* landing site was mutated using CRISPR (Cho *et al.* 2013; Paix *et al.* 2015). Three Unc-119 lines were obtained from individuals injected with Cas9 protein and 3 *unc-119* sgRNAs. One allele had a 691 bp deletion *(Cb-unc-119(ant40)).* This mutated landing site transgene was named *antIs31* and then backcrossed to N2 4 times, each time retaining Unc-119 individuals, to yield BRC0566, the main landing site strain used for injections.

### Recombination mediated cassette exchange (RMCE) plasmids

Plasmid names, injection concentrations, and descriptions are in Sup Table S1. For the initial proof-of-principle RMCE experiments plasmid, pBRC_double_attB_GFP, containing *sur-5p::gfp* flanked by attB sites was constructed using conventional cloning. To facilitate the RMCE experiments using the Unc-119 strain (BRC0566), two general plasmids were constructed. The first plasmid, pBRC_double_attB_GFP_donor, contains a GFP marker to facilitate visually following the insertion; it carries oppositely oriented attB sites flanking *Cb-unc-119(+), sur-5p::gfp* and the R1-R2 Gateway cloning sites. The second plasmid, pCG150_double_phiC31_attB, lacks fluorescent markers and carries oppositely oriented attB sites flanking *Ce-unc-119(+)* and the R3-R4 Gateway cloning sites. Conventional restriction site mediated cloning is also possible for both plasmids. Plasmid pBRC_double_attB_GFP_mCherry was generated by Gateway cloning into pBRC_double_attB_GFP_donor. The operon reporters carrying two fluorescence colors were constructed by conventional cloning into pCG150_double_phiC31_attB.

For the BAC insertion experiment, the fire ant BAC 47D10 was retrofitted with a cassette containing two attB sites, *Cb-unc-119(+),* and the kanamycin resistance gene to yield attB_retrofitted_Si47D10 using recombineering (Quick & Easy BAC Modification Kit, Gene Bridges). Plasmids were isolated using the Mini Plus Plasmid DNA Extraction System (Viogene, cat no. GF2002); the BAC was isolate using the Qiagen Genomic-tip 20/G (Qiagen, cat no. 10223).

### Recombination mediated cassette exchange experiments

For the initial landing pad strain BRC0546 *(antIs30),* adult (non-Unc) hermaphrodites were injected using standard methods (Evans (ed.) 2006) with the donor cassette *(sur-5p::gfp)* and the co-injection marker *(myo-2p::RFP)* plasmids. Then, transgenic F1 were singled out based on GFP fluorescence. After, F2 and subsequent progeny were then screened for Unc, Green, and non-Red individuals, who were putatively homozygous for the *sur-5p::gfp* RMCE insert.

For the improved landing pad strain BRC0566 *(antIs31),* adult (Unc) hermaphrodites were injected as above. Then, transgenic non-Unc F1 were singled out. Next, F2 and subsequent progeny were screened for non-Unc individuals that segregated all non-Unc individuals, and hence were putatively homozygous lines. Screening for homozygous lines was attempted up to the F6 generation if approximately three-quarters of the progeny were non-Unc at each generation. Otherwise, the transgene was presumed to be an extrachromosomal array and screening was stopped. For candidate homozygous lines, the GFP and mCherry fluorescence patterns were used to help ascertain if the RMCE insertion was likely correct. Specifically, there should be a loss of the *myo-2p::gfp* within the landing pad and the co-injection marker mCherry as well as gain of any donor construct GFPs and/or mCherry.

PCR and sequencing assays were conducted to validate the RMCE insertions for all candidate correct insertion strains. First, a set of four PCR assays using two flanking the landing pad and two construct-specific primers near the edge of insert were used to confirm both integration and orientation. Positive PCR products were also sequenced to verify that recombination produced two proper attR sites. Second, two PCR assays were used to test if the *Cb-unc-119* and phiC31 integrase genes were lost. Third, for the dual cassette experiment, a series of PCR assays were designed spanning various subparts across the whole insert. Finally, in some cases, long range PCR assay across the entire insert was conducted. Primers are in Sup Table S2.

### RMCE with a BAC

The attB_retrofitted_Si47D10 BAC was injected, as for the plasmids above, in approximately half of the injections; in the remaining injections spermidine and spermine (70 μM and 30 μM, respectively) was added and the co-injection marker was omitted. Screening for BAC integrants also included PCR assays for the gain of the bacterial kanamycin resistance gene, the loss of *myo-2p::gfp,* and seven parts across the BAC spaced about 10-15 kb apart. All PCR assays were also conducted on the known incorrect BAC insertion lines, i.e., those with GFP and/or mCherry expression. Southern blot analysis was conducted according to standard protocol with random primer labeling using ^32^P-dCTP. Primers used for the BAC validation assays are in Sup Table S2.

### Dual color operon reporter experiments

Three operon reporters carrying two fluorescence colors were injected into BRC0566 to obtain single copy integrants. The seam cell specific operon reporter plasmids, SCMp::GFP(PEST)::H2B::lin-41 3′UTR::SL2::mCherry::H2B::unc-54 3′UTR, contains the seam cell specific promoter expression an operon containing GFP and mCherry, both as histone chimeras. GFP is also fused to a PEST degron and contains the *lin-41* 3′-UTR or *lin-41* 3′UTRΔLCS (LCS, let-7 complementary sequence) (Rausch *et al.* 2015). An SL2 sequence, containing the *gpd-2/gpd-3* intergenic region (Merritt *et al.* 2008), precedes mCherry which contains the *unc-54* 3′-UTR. The ubiquitously expressed reporter plasmid, dpy-30p::GFP(PEST)::H2B::lin-41 3′UTR::SL2::mCherry::H2B::unc-54 3′UTR, was similarly constructed except with the *dpy-30* promoter (Sup. Fig. S1). These operon reporters were cloned into pCG150_double_phiC31_attB (Sup. Fig. S2) for injection into BRC0566 with co-injection marker egl-20p::mCherry. The seam cell specific construct was also crossed into the *let-7(n2853)* hypomorphic mutant to test the post-transcriptional regulatory role of *let-7* on the *lin-41* 3′-UTR. For validation, we conducted a double ‘PCR walking’ assay: one primer was fixed to the left or right flanking region of *ttTi5605* and others primers were staggered approximately every 1.5 kb along the transgene (Sup. Fig. S3). After validation of the insertion lines, images were taken of the worms and then GFP and mCherry expression levels were quantified.

For microscopy assays, synchronized arrested L1 larvae of animals carrying dual-color reporter in wildtype or *let-7(n2853)* background, were plated on food and incubated at 20°C on NGM agar plates with *Escherichia coli* OP50 bacteria and imaged at the L2 and mid L4 stage (18 and 40 h after plating). Images were acquired in the green, red and transmitted light channels (with Differential Interference Contrast, DIC) using a 63× Zeiss oil immersion objective on a Zeiss fluorescence microscope (Axio Imager.M2) and Axiovision 4.8 software. For data analysis, worms were selected based on visual inspection of gonad length and vulva morphology (Mok *et al.* 2015). Seam cells were selected by Fiji. Images were denoised using a Gaussian blurred algorithm and segmented using a Yen global threshold. Signal intensity in the green channel was divided by that in the red signal for each cell; relative signal intensities were then averaged for each worm. Two to 10 seam cells in 11-14 worms per genotype were quantified, medium signal intensity and frequency of the signal intensity were calculated and graphed using GraphPad Prism software.

### Generation of chromosomal inversion on LG IV

co-CRISPR (Arribere *et al.* 2014; Paix *et al.* 2015) was used to generate one strain carrying the minimal 30 bp attP site inserted into the second intron of *dpy-13* and another carrying a 38 bp attB site in the fourth intron of *unc-30.* The sgRNAs, repair templates, and PCR primers used for validation are listed in Sup Table S2. The insertion alleles obtained were *antSi50* (attB, BRC0609) in *dpy-13* and *antSi51* (attP, BRC0699) in *unc-30;* these strains are wild-type (WT) and have been backcrossed 3x to N2. Recombination was used to place the attB and attP insertions onto the same chromosome to generate BRC0671 *(antSi50 antSi51)*.

The chromosomal inversion was generated as follows. First BRC0566 *(antIs31; unc-119(ed9))* was outcrossed to N2 to remove *unc-119(ed9)* to yield BRC0664. Next, N2 males were crossed to BRC0664 and then heterozygous F1 males were crossed to BRC0671 to produce *antIs31/+; antSi50 antSi51/+ +* or *+/+; antSi50 antSi51/+ +* F2 cross progeny. *myo-2p::gfp* is difficult to detect in adults, so it was scored in the F3 progeny. F3 and F4 progeny were screened for Dpy Unc individuals, indicative of an inversion mediated by phiC31 integrase. Inversions were confirmed in the independently derived Dpy Unc lines by PCR and sequencing of the junctions.

Recombination rates between *dpy-13* and *unc-30* in the inversion allele, *antIn1,* and in the wild-type orientation *(dpy-13(e184) unc-30(ok613))* were determined from the recombinant offspring of the *dpy-13 unc-30/+ +* heterozygotes. The recombination rate was calculated based on the fraction of non-Dpy non-Unc F2 that segregated non-Dpy Unc or Dpy non-Unc and the Punnett Square expectations for recombinants.

### Data availability

The authors state that all data necessary for confirming the conclusions presented in the article are represented fully within the article. All data generated is present in figures, supplemental figures, tables and supplemental tables. Critical worm strains and plasmids will be made available at the CGC and Addgene, respectively. All other reagents are available upon request from Shih-Peng Chan and John Wang.

## RESULTS

### RMCE proof of principle

To adapt the phiC31 recombination system for *C. elegans*, we created a strain possessing a landing site for RMCE consisting of inverted attP sites that surround a GFP marker and a germline expressed phiC31 integrase (Fig. 1A). This strategy was chosen for the following reasons. First, injections of the desired sequence (or construct) for RMCE would be simplified, because co-injection of phiC31 integrase (as DNA, mRNA, or protein) would not be needed. Second, because phiC31 integrase protein would already be present, integration may be more efficient. Third, by placing the phiC31 integrase between the attP sites, RMCE would swap out the phiC31 integrase from the chromosome, eliminating the need to conduct crosses to remove the integrase. Finally, the GFP marker would permit visually tracking the presence of the landing site in crosses or its absence after RMCE.

**Figure 1.**
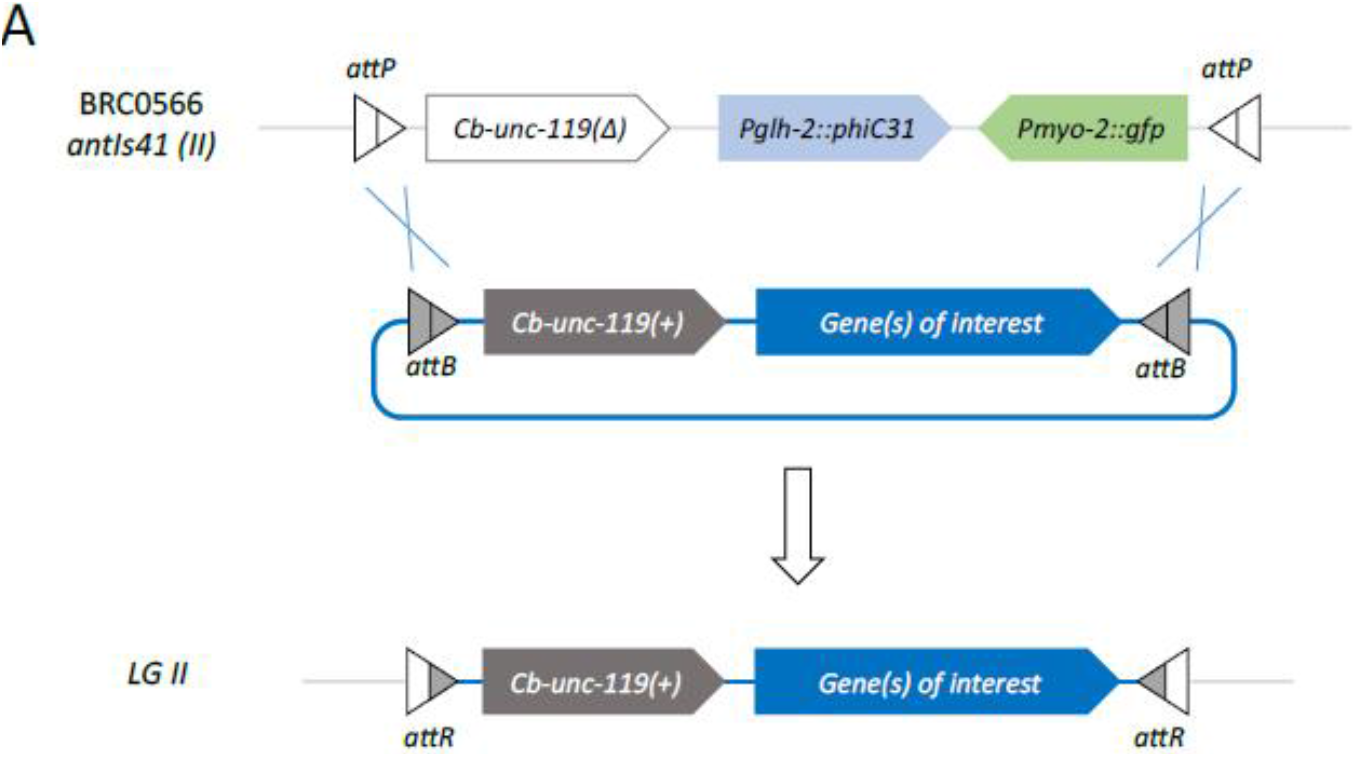
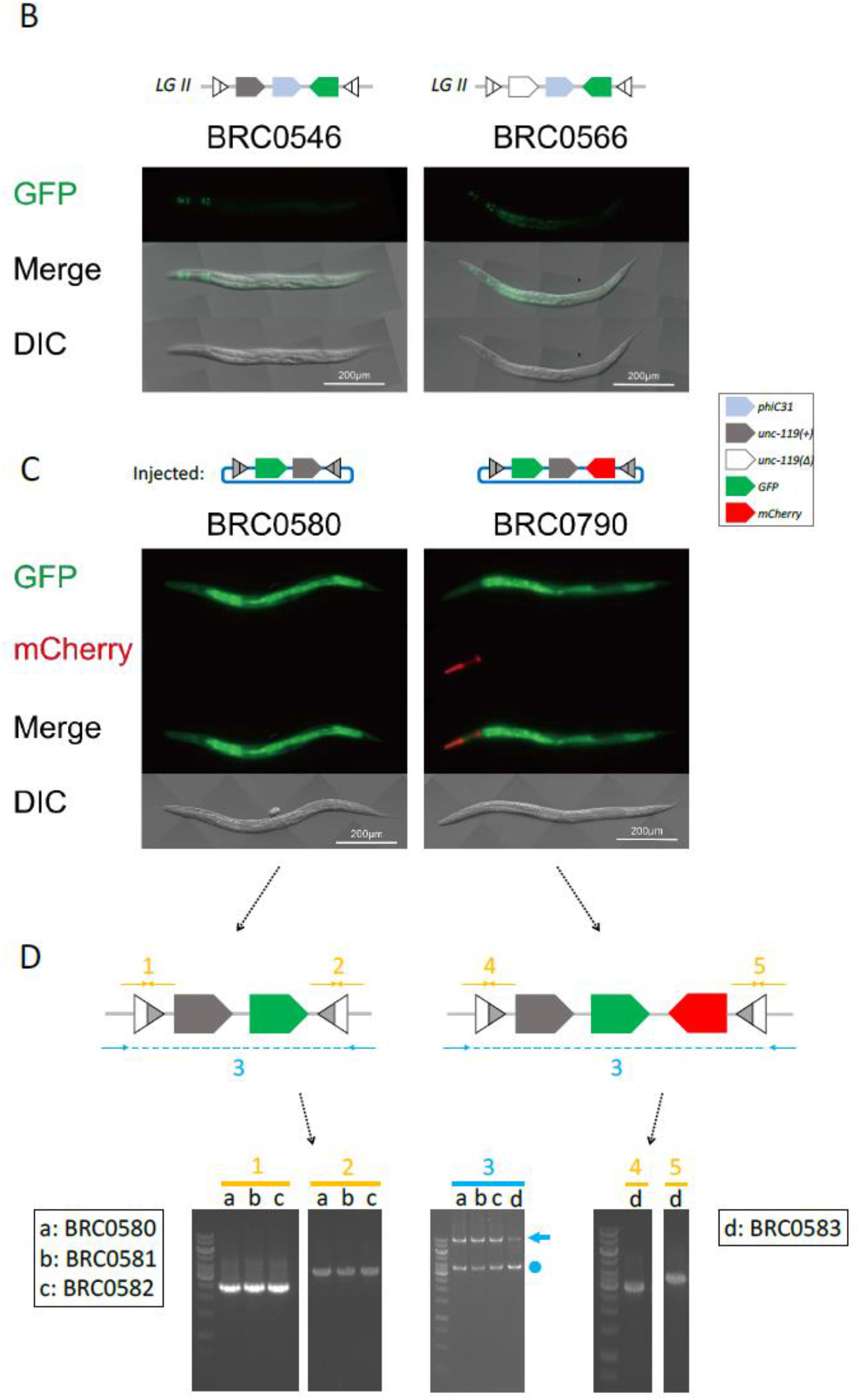
phiC31 integrase mediated RMCE in *C. elegans.* (A) Schematic of single copy transgene integration via RMCE into the *antIs41* docking site. (B) Strains used for RMCE. Both BRC0546 and BRC0566 express phiC31 integrase in the germline. BRC0566 is derived from BRC0546 after mutating the *Cb-unc-119* gene. GFP expression in the pharynx from the *myo-2* promoter is evident; weak expression in other parts of the body is also seen. (C) Images of two representative strains after successful RMCE. BRC0580 carries a single copy integrant of *sur-5p::gfp;* BRC0790 carries a single copy integrant of *sur-5p::gfp* and *myo-2p::mCherry.* Locations of GFP and mCherry expression match the expected pattern. (D) Validation of proper integrations with PCR assays for four representative strains. Orange PCR assays indicate test for proper recombination junctions and orientation. Integrants can also be in the reverse orientation, and those PCR assays are not shown. Blue indicates long PCR assays across the entire insert and demonstrate single copy insertion. Arrows next to the gels indicate the correct long PCR product; dots are PCR artifacts caused by probable intramolecular annealing of the attR sites.

We generated this phiC31 integrase expressing landing site strain using the MosSCI technique to target a construct containing the desired elements and *Cb-unc-119(+)* into the *ttTi5605* Mos1 insertion site on LG II (Frokjaer-jensen *et al.* 2008). This site was chosen because it is permissive for germline expression of transgenes (Frokjaer-jensen *et al.* 2008). After screening the progeny from approximately 10,000 young adult worms subjected to MosSCI, we succeeded in obtaining one correct integrant (Fig. 1B, BRC0546; *antIs30 [attP-phiC31-f Cb-unc-119(+) glh-2p::phiC31 myo-2p::GFP attP-phiC31-r]; unc-119(ed9)).* We further verified that phiC31 integrase was expressed and properly spliced (data not shown).

Next, we tested whether the *antIs30* phiC31 integrase expressing landing site could be used for RMCE. We created a donor plasmid containing a globally expressed green fluorescent protein gene *(sur-5p::GFP)* that is flanked with inverted attB sites; this sequence to be inserted is 2,559 bp. Recombination between both pairs of attP/attB sites would swap in *sur-5p::GFP* and swap out phiC31 integrase, *Cb-unc-119(+),* and *myo-2p::GFP.* Compared to the original strain, homozygous recombinant individuals would have a switch in GFP expression location from the pharynx to the whole body and also from wild-type (WT) to the Unc phenotype.

We injected this donor plasmid and a co-injection marker *(myo-2p::mCherry)* into 26 *antIs30; unc-119(ed9)* P0s and then singled out all 28 WT F1 individuals that were transgenic for the donor plasmid based on their expression of *sur-5p::GFP.* Three F1s continued segregating F2 individuals expressing *sur-5p::GFP* (Table 1). In one case the F2 brood was composed of mostly WT animals and many Unc animals putatively homozygous for the donor construct and lacking mCherry expression (i.e., Unc, Sur-5p::GFP, and non-Myo2p::GFP), consistent with the F1 mother being heterozygous for the donor construct. In the second case, all the F2 were WT, with some expressing *sur-5p::gfp* and others not. We singled out nine *sur-5p::gfp* F2 individuals and found that all segregated many Unc, Sur-5p::GFP, and non-Myo2p::GFP F3 progeny, indicating that these F2 were likely heterozygous for the donor construct. The last F1 individual segregated F2 and subsequent F3 in a pattern that appeared consistent with only an extrachromosomal array (i.e., no integration) and was not examined further.

**Table 1.** Summary of phiC31-mediated RMCE experiments

| Experiment<sup>a</sup> | P0 injected | Transgenic F1 | Integrated lines | Correct RMCE lines<sup>b</sup> | % Correct lines |
| --- | --- | --- | --- | --- | --- |
| pBRC_double_attB_GFP<sup>c,d</sup> | 26 | 28 | 2 | 2 | 100% |
| pBRC_double_attB_GFP_donor<sup>d</sup> | 165 | 104 | 7 | 5 | 71% |
| pBRC_double_attB_GFP_mCherry<sup>d</sup> (total) | 238 | 207 | 33 | 25 | 76% |
| trial 1 | 180 | 80 | 5 | 3 | 60% |
| trial 2 | 47 | 112 | 25 | 19 | 76% |
| trial 3 | 11 | 15 | 3 | 3 | 100% |
| SCMp::GFP(PEST)::H2B::lin-41 3'UTR::<br>SL2::mCherry::H2B::unc-54 3'UTR<sup>e</sup> | nd | >20<sup>f</sup> | >6<sup>g</sup> | 6<sup>h</sup> | >30% |
| SCMp::GFP(PEST)::H2B::lin-41 3'UTR mut::<br>SL2::mCherry::H2B::unc-54 3'UTR<sup>e</sup> | nd | >20<sup>f</sup> | >6<sup>g</sup> | 6<sup>h</sup> | >30% |
| dpy-30p::GFP(PEST)::H2B::lin-41 3'UTR::<br>SL2::mCherry::H2B::unc-54 3'UTR<sup>e</sup> | nd | >20<sup>f</sup> | >6<sup>g</sup> | 6<sup>h</sup> | >30% |

<sup>a</sup>Plasmids injected into BRC0566(*antIs31*) resulting in rescue of *unc-119* mutants into wild-type, unless indicated
<sup>b</sup>Based on expected GFP and mCherry phenotypes as well as PCR for the recombination junctions
<sup>c</sup>In this experiment, the plasmid was injected into BRC0546(*antIs31*) and homozygous RMCE integrants would have an Unc phenotype
<sup>d</sup>Co-injection plasmid was *myo-3p*::mCherry
<sup>e</sup>Reporter construct was cloned into pCG150\_double\_phiC31\_attB before injection; Co-injection plasmid was *egl-20p*::mCherry
<sup>f</sup>Only 20 F1s were picked although more present
<sup>g,h</sup>Only the first 6 phenotypically correct integrated lines were examined by PCR assays and all 6 had the correct recombination junctions nd, not determined

To confirm that the Unc worms were the product of phiC31 mediated RMCE at the landing site, we conducted molecular validation experiments on the two independent strains. PCR assays for the two recombination junctions, for the presence of phiC31 integrase, and with flanking primers across the landing site were consistent with the donor *sur-5p::gfp* construct replacing the *phiC31* integrase at the landing site (data not shown). Additionally, sequencing of the junction PCR products verified that both recombination junctions were attR sites, the outcome of attP x attB recombination. Together these results demonstrate that phiC31 mediated RMCE is possible in *C. elegans,* can occur at relatively high frequency (2 out of 3 transgenic lines), and integration can occur in the F1 or the F2 generations.

### Improving and facilitating RMCE

Although the strain BRC0546 *(antIs30; unc-119(ed9))* can be used for RMCE, identifying recombinants is demanding because it requires screening for Unc individuals in the context of WT individuals. Screening for WT individuals rescued for the Unc phenotype would be easier. Therefore, we used CRISPR/Cas9 to mutate the *Cbr-unc-119* gene within the landing site, yielding *antIs31[attp-phiC31-f; Cbr-unc-119(ant40); Pglh-2::phiC31; Pmyo2::GFP; attp-phiC31-r]* (Fig. 1B). We then backcrossed this strain four times to N2 to produce BRC0566 *(antIs31 ant40* II*; unc-119(ed9)* III).

To facilitate generating donor constructs for RMCE, we developed two plasmids (Sup. Table S1). The first one (pBRC_double_attB_GFP_donor) has *sur-5p::GFP,* which permits visual following of the recombinant transgene in subsequent crosses, while the second one (pCG150_double_phiC31_attB) does not, which permits experiments where tracking of custom fluorescent signals is desired. For both plasmids, the gene of interest can be added into these plasmids by standard or Gateway cloning. Finally, both plasmids have *Cbr-unc-119(+)* for screening.

We next used these plasmids to confirm that RMCE in BRC0566 works (Table 1). We injected the pBRC_double_attB_GFP_donor plasmid, which would insert ~6.4 kb at the landing site, along with an mCherry co-injection marker into BRC0566. We obtained 104 transgenic F1 that gave rise to seven putatively integrated lines having the predicted recombinant phenotype (WT, Sur-5p::GFP, and non-Cherry, Fig. 1C). Of these, five (71%) were validated to be correct (Fig. 1D). Because the pairs of attP sites in the genome and of attB sites on the donor construct are identical, the donor construct can integrate in two orientations, which we call sense and anti-sense. We obtained two and three such lines, respectively. We also injected a plasmid where Gateway cloning was used to insert *myo-2p::mCherry* into pBRC_double_attB_GFP_donor, creating a construct with an insert size of ~7.4 kb. In a series of three experiments, we obtained 25 successfully integrated lines with success rates ranging from 60% to 100% (Fig. 1C, D, Table 1). Finally, we used conventional cloning to insert various two gene operons into pCG150_double_phiC31_attB to create donor inserts of ~9 to 15 kb. These operon constructs were designed as a proof-of-concept platform for dual expression (more below). Again, we easily obtained proper insertions that were molecularly validated (Fig. 2, Table 1, and Sup. Figs. S1-3). Together these results demonstrated that phiC31-mediated RCME can be used in this improved strain and that larger insertions up to ~15 kb can be generated relatively easily.

**Figure 2.**
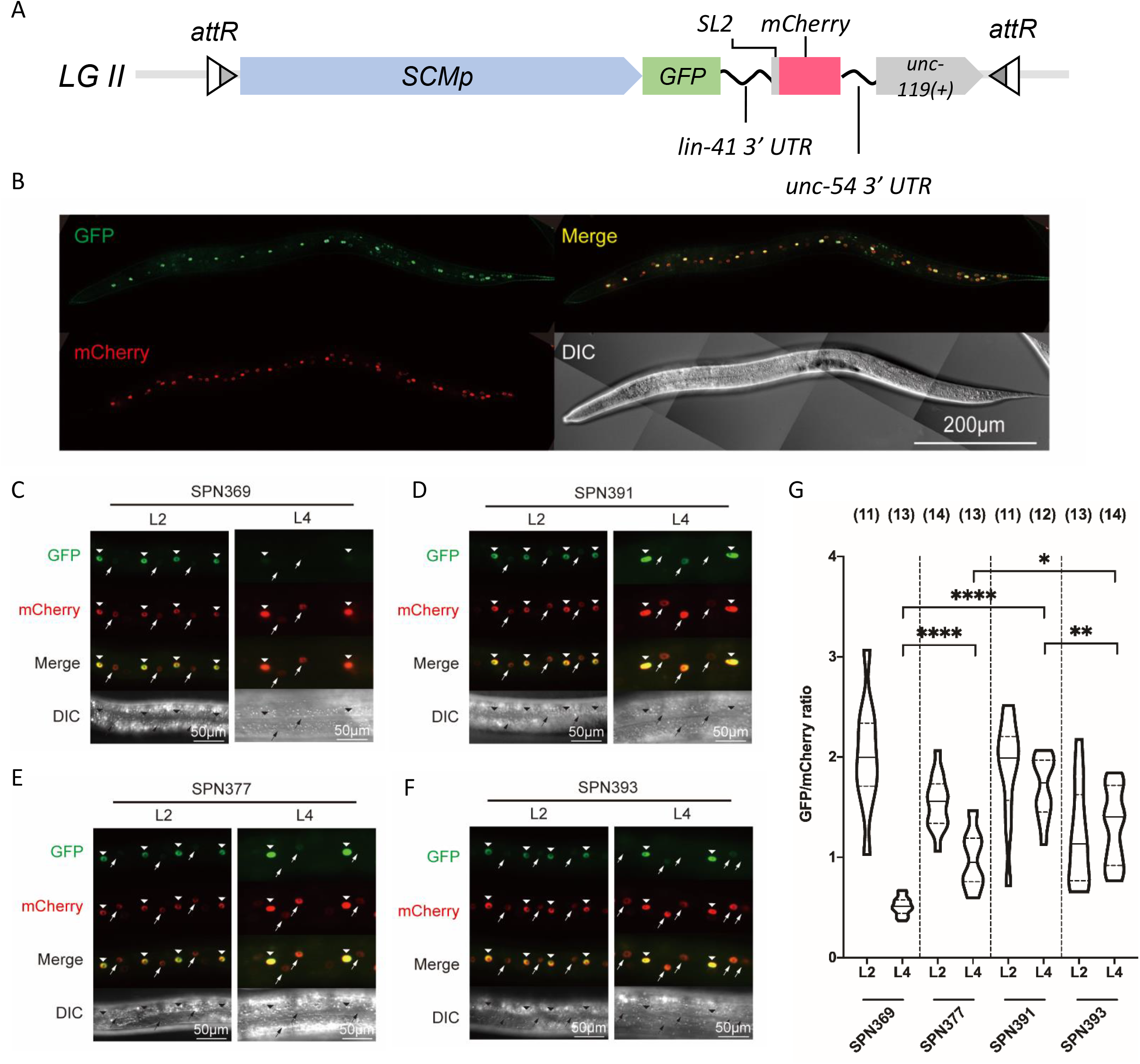
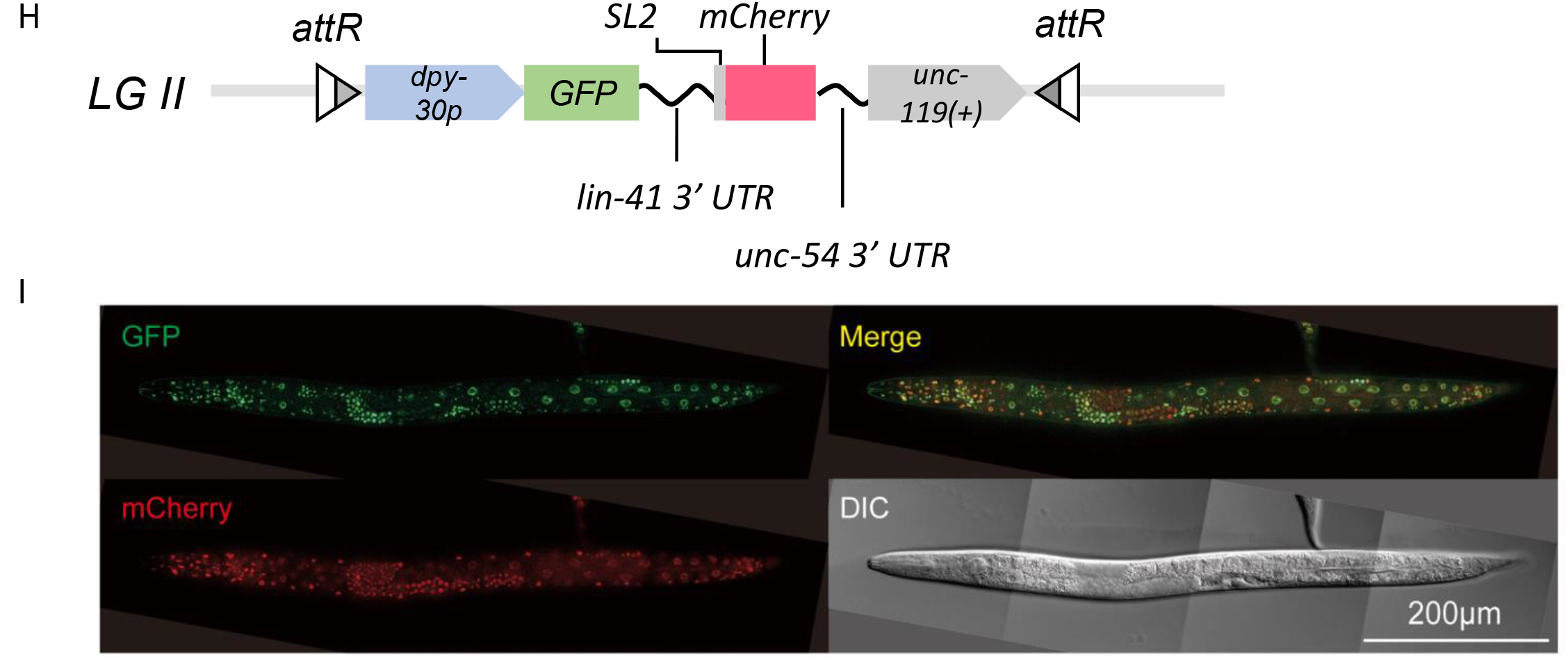
Measuring the activity of *let-7* using a dual-color reporter. (A,H) Schematic of the dual-color reporter used to monitor miRNA activity in *C. elegans.* Transcription of the single-copy integrated reporter is driven by (A) a seam cell specific promoter or (H) the ubiquitously active *dpy-30* promoter. The GFP reporter fused to the *lin-41* 3′ UTR contains two *let-7* binding sites (as shown in Sup. Fig. S1) that are regulated by the *let-7* miRNA, while mCherry fused to the *unc-54* 3′ UTR serves as an internal control. (B,I) The expression of dual-color reporter with (B) a seam cell specific promoter or (I) the *dpy-30* promoter in the mid L4 stage in wild-type *C. elegans.* (C,D) Repression of a *let-7* target reporter *(chsIs001)* in mid L4 worms depends on *let-7*(+) (SPN369) and is lost *let-7(n2853)* mutants (SPN391). (E,F) *lin-41* 3′ UTR reporter without the *let-7* binding sites *(chsIs002)* in mid L4 worms is regulated neither in *let-7* wild-type (SPN377) nor in *let-7(n2853)* mutants (SPN393) individuals. (C,D,E,F) L2 staged animals serve as *let-7* negative controls. Seam cells are marked with arrowheads and hyp7 nuclei are marked with arrows. (G) Quantification of reporter expression in worms. Violin plots show the GFP signal intensity divided by the mCherry intensity for each worm per condition. Two to 10 seam cells were quantified per worm. The number of worms examined for each condition are indicated on top. Horizontal line and dotted lines indicate median value and first/third quartiles, respectively. \**P* < 0.05 and \*\*\*\**P* < 0.0001, two-tailed unpaired t-test.

### Single copy dual-fluorescent operon reporter system

A two-color fluorescent reporter system has been developed to more accurately quantify post transcriptional control by microRNAs (miRNA) at the single cell level (Ecsedi *et al.* 2015). In this system two reporters are driven by the same promoter but have different 3′-UTRs. A test 3′-UTR is fused to GFP while a control, unregulated 3′-UTR is fused to mCherry. Thus, regulation of the 3′-UTR such as via miRNAs should be reflected by changes in the GFP/mCherry signal ratio. To maintain robustness of the signal, the GFP and mCherry reporters are integrated site-specifically in single copy. However, because of insertion size limitations (by MosSCI or CRISPR/Cas9) two integration events, onto separate chromosomes, are required.

We took advantage of the larger cargo size of the phiC31 integrase system to streamline the dual fluorescent system and put both reporters at one genomic location. We designed an operon reporter that permits examining the heterochronic downregulation of the *lin-41* 3′UTR by the *let-7* miRNA in the seam cells using quantitative fluorescence microscopy. This operon reporter consisted of a seam cell specific promoter (SCMp) driving *GFP(PEST)::H2B::lin-41 3′UTR* and *mCherry::H2B::unc-54 3′UTR* (Sup. Fig. S1). The *gpd-2/gpd-3* intergenic region connects the two markers and provides a SL2-specific trans-splice site for the *mCherry* (Huang *et al.* 2001). We cloned this operon reporter into plasmid pCG150_double_phiC31_attB (constructed in this study, Sup. Fig. S2), injected the donor construct into the Unc landing pad strain (BRC0566), and obtained at least 6 WT candidate integrants. One of them was verified for proper RMCE insertions by double ‘PCR walking’ (see methods, Sup Fig. S3A).

We next used one of the integrated alleles, *chsIs001,* to establish that this operon correctly reports let-7/LIN-41 3′-UTR regulation. *let-7* is upregulated during the L4 stage. Thus, we followed GFP/mCherry expression ratios during development with the expectation that the expression ratio at the L4 stage should be less than at the L2 stage. As predicted, we observed reduced GFP/mCherry expression ratios in the seam cells (Fig. 2C). To confirm that *let-7* is necessary for the decrease in GFP expression, we crossed the dual reporter strain into the *let-7(n2853)* hypomorphic mutant and found that the GFP signal at the L4 stage were not down-regulated as in wild-type (Fig. 2D). Finally, to test the requirement of the *let-7* complementary target sequences on the *lin-41* 3′-UTR, we generated a new phiC31-mediated RMCE operon integrant *(chsIs002)* with the target site deleted. Examination of one strain revealed no reduction of the GFP/mCherry ratio at the L4 stage in both the WT or *let-7(n2853)* backgrounds (Fig. 2E and 2F). Quantitation of the GFP/mCherry ratios are shown in Fig. 2G.

To expand the utility of the reporter to the whole animal, we replaced the SCM promoter with a ubiquitously expressing *dpy-30* promoter (Fig. 2H). We used phiC31-mediated RMCE of this construct to obtain worm lines stably expressing GFP and mCherry in almost all tissues, including the gonad *(chsIs004,* Fig. 2I and Sup Fig. S3C). Interestingly, the GFP/mCherry expression ratio varied along the gonadal axis, likely reflecting spatial-temporal regulation of the LIN-41 (or possibly of the UNC-54) 3′-UTR in this organ. In addition, the GFP/mCherry expression ratio differed among cell types, and was easily noticed in the intestines and some neuron cells, further suggesting cell-type regulation of the LIN-41 3′-UTR.

### BAC integration

The ability to generate large single copy transgenes (10s to 100+ kb) at specific locations would facilitate more complex experiments. To this end we tested whether it would be possible to integrate a bacterial artificial chromosome (BAC). We used recombineering to retrofit an ~130 kb fire ant BAC (clone 47D10) with a cassette that contains *Cb-unc-119(+), Psur-5::GFP,* and two attB sites. The attB sites are oriented such that the entire BAC (including vector sequence), except for *Psur-5::GFP,* would be inserted upon proper RMCE (Sup. Fig. 4A).

We injected this retrofitted BAC both with and without an *mCherry* co-injection marker into the landing site strain and obtained 138 F1 non-Unc individuals. Of these, 24 gave rise to putatively integrated lines based on segregating exclusively non-Unc progeny. Based on PCR assays nine of these had proper junctions consistent with phiC31 mediated recombination and had lost phiC31. Nevertheless, seven of the lines still expressed myo-2p::GFP, sur-5p::GFP, and/or mCherry indicating that clean cassette exchange did not occur and suggesting that extrachromosomal array formation may have occurred prior to phiC31 mediated integration. We did obtain 2 lines that expressed neither GFP nor mCherry, which would be expected of potential RMCE integrants. However, neither were proper, clean single copy insertions based on additional PCR assays and Southern analysis (Sup. Fig. 4B and 4C). Together this indicates that inserting single copy BAC-sized constructs through phiC31 mediated recombination in *C. elegans* is not readily possible with our injection conditions but may still be possible with technical improvements.

### Generating large inversions

phiC31 integrase can be used to invert sequences that are flanked by oppositely oriented attP and attB sites (Merrick *et al.* 2018). To test if it is possible to generate a very large inversion in *C. elegans* we first created a strain carrying the attP and attB sites in opposing orientations ~9 Mb apart on LG IV. The attP *(antSi50)* and attB *(antSi51)* sites were inserted into an intron of the *dpy-13* and *unc-30* genes, respectively (Fig. 3A). Because the insertions are in introns, both insertion alleles are WT. However, an inversion would truncate both genes and produce a Dpy Unc phenotype.

**Figure 3.**
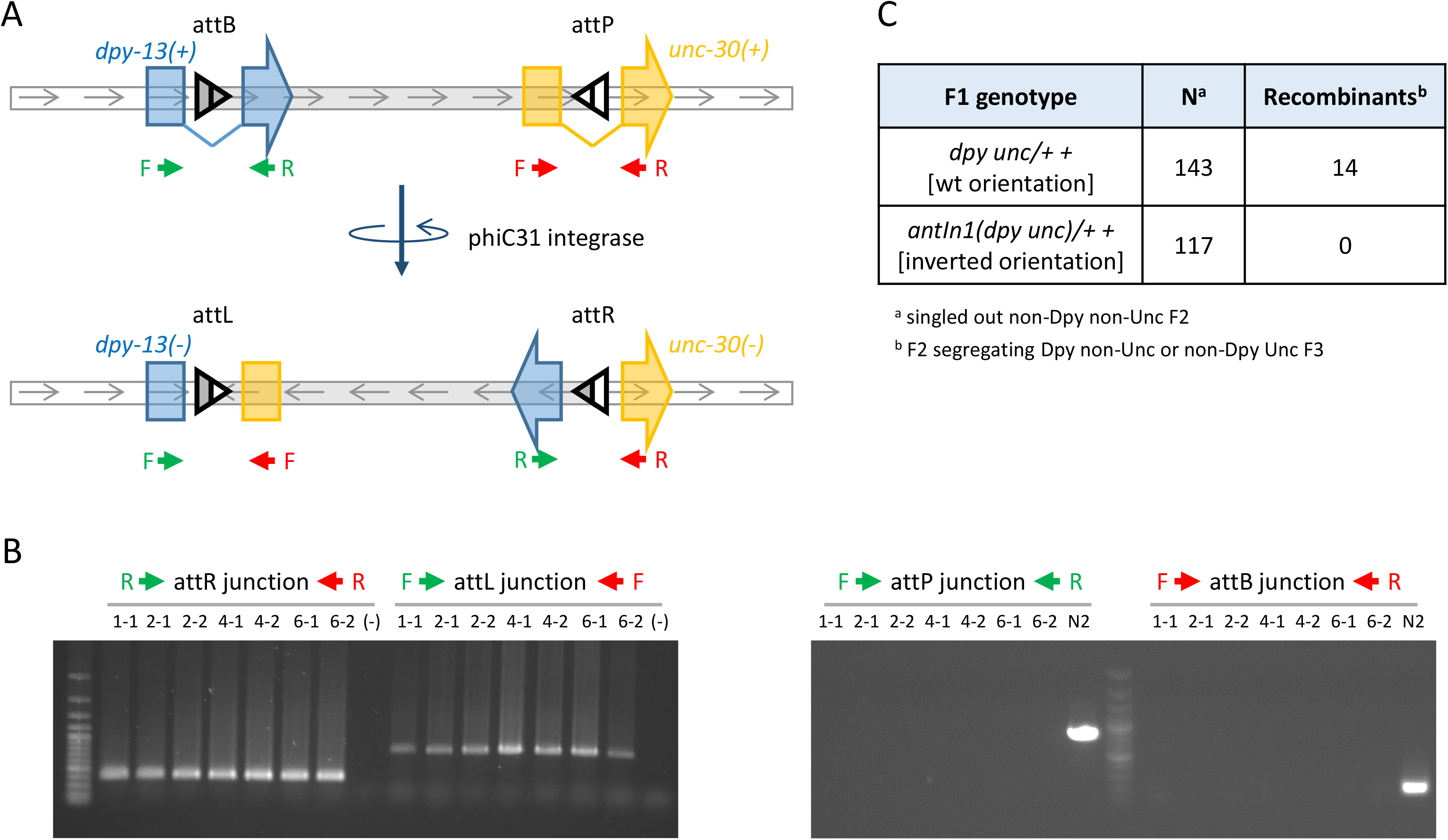
Generation of a large balancer inversion. (A) Schematic for generating an inversion spanning the *dpy-13* to *unc-30* interval on LG IV. An attB and an attP site were inserted into an intron of the *dpy-13* and *unc-30* genes, as indicated. Worms carrying this chromosome were crossed to a phiC31 integrase expressing strain and then Dpy Unc strains carrying homozygous inversions were isolated. Red and green arrows represent primers used in PCR assays testing for the original or inverted orientations. Intron and exon structure are simplified and are illustrative only; gene sizes and distances are not to scale. (B) PCR assays for representative Dpy Unc strains that carry the phiC31 integrase induced inversion. Seven strains from four independent lines (first digit of lane IDs) are shown. PCR arrays for the recombinant junctions are positive while those for the original orientations are negative, except for N2. Minus sign (-) is water. (C) The inversion, *antIn1,* suppresses recombination in the *dpy-13* to *unc-30* interval.

To induce the inversion, we crossed the phiC31 integrase expressing transgene, *antIs31,* into the attP and attB bearing strain to generate *antIs31/+; antSi50 antSi51/+ +* heterozygotes. From 15 F1 heterozygotes, we obtained 8 independent Dpy Unc lines, via individuals found in the F2 or F3 generations. We then used PCR assays to show that the Dpy and Unc phenotypes were consistent with an inversion within the *dpy-13* and *unc-30* genes (Fig. 3B). This result demonstrates that phiC31 integrase can mediate precise inversions in *C. elegans* and such inversions can be multi-megabase in size.

Inversions should suppress recombination. To determine the recombination rate in the *dpy-13 unc-30* interval for the phiC31 mediated inversion, we crossed N2 males to one of the inversion alleles, *(antIn1[dpy-13(ant41) unc-30(ant42)]),* obtained F1 heterozygotes *(dpy-13 unc-30/+* +), singled out non-Dpy non-Unc F2 individuals, and then scored for F2 heterozygous recombinants (reflecting recombination events in the F1 mother) by examining the F3 segregation pattern. None of the 117 singled out F2 individuals segregated any recombinant genotypes (Fig. 3C); this proportion is different from that expected for the 8.22 cM genetic distance between the two genes (*P* = 2.0 x 10e-6, expect 13.0, exact binomial test). Control crosses using the double mutant *dpy-13*(e184) *unc-30(ok613)* that is in the wildtype orientation yielded 14/143 recombinants (Fig. 3C), which corresponded to 7.2 cM, and was not different from expected (*P* = 0.79, expect 15.9, exact binomial test). These results demonstrate that *antIn1* strongly suppresses recombination in the *dpy-13 unc-30* interval

## DISCUSSION

We have demonstrated that the phiC31 integrase system can be used in *C. elegans* to insert transgenes through RMCE. RMCE insertion rates are reasonably high for insertions up to ~15 kb (~60% on average). Using phiC31 mediated RMCE, we developed a system to study post-transcriptional gene regulation by integration of a dual fluorescence operon reporter. Additionally, we showed that phiC31 integrase can be used to generate inversions, and such inversions can be large.

phiC31 integrase RMCE provides another tool to complement MosSCI, CRISPR/Cas9, and Flp/FRT for precise single copy insertion of transgenes in *C. elegans.* The main advantages of our current system are the larger size of insertion allowed and simple protocol. With respect to insert size, the Flp system has comparable efficiencies for obtaining insertion sizes in the 10-20 kb range, but MosSCI and CRISPR/Cas9 are more laborious for insertion sizes greater than ~2 kb (Frokjaer-jensen *et al.* 2008; Dickinson and goldstein 2016). Biolistic bombardment (Praitis *et al.* 2001) and miniMos (Frokjaer-jensen *et al.* 2014) are alternatives for generating single or low copy insertions, but insertion sites for both methods are random. Our current protocol is also fairly straightforward, requiring only the injection of two plasmids, a donor construct and a co-injection marker, followed by screening for WT integrants. Additionally, no extra manipulations or crosses are required to remove the phiC31 integrase. In comparison, the CRISPR/Cas9 and Flp systems typically requires cloning out rollers and then an additional self-excision step to remove the roller and/or other positive selection markers (Nonet 2020). Direct integration of short inserts with MosSCI can be comparably simple, but larger insertions require generating an extrachromosomal array first followed by a laborious screening step. The efficiency and simplicity of our system also make obtaining insertion strains fairly fast. We have occasionally obtained homozygous individuals already in the F2 generation, and almost always by the F3. Overall this permits rapid tests of genetic constructs especially when single copy is important.

Another advantage is that phiC31 integrase mediates unidirectional recombination between attP and attB sites, which are dissimilar, to generate attR and attL sites, which are also dissimilar. Thus, insertions cannot be excised. In contrast, in the Flp based system, the FRT recombination sites are the same before and after recombination, so an insertion could be subject to immediate excision. Unidirectional recombination also permits additional rounds of phiC31 RMCE into strains already containing RMCE transgenes. The Flp system may result in the excision of prior integrated transgenes.

Our current design does have several limitations. First, we currently have only one landing site for integration. Additional landing sites on different chromosomes would offer more experimental flexibility, especially for conducting downstream crosses or for testing different local genomic environments. Second, integrations into the landing site can occur in two orientations because the two attP sites are identical in sequence. This could be resolved by placing different attP variants on the “left” and “right” end of the landing site (with matching changes for the attB sites on the donor construct) to force insertion in a specific orientation, like has been done for the Flp/FRT system (Dickinson *et al.* 2015; Nonet 2020). A third issue is that the effect of constitutive germline expression of phiC31 integrase is unknown. Constitutive expression of phiC31 integrase in the *D. melanogaster* germline seems to have little deleterious effect (Bateman *et al.* 2006). Similarly, *antIs31* does not exhibit any overt negative phenotypes. Nevertheless, we did lose the transgene once after passaging for over 1 year. Thus, continuous expression of phiC31 integrase might be slightly deleterious. Another potential shortcoming is that the donor construct contains *unc-119(+),* which may not be desirable. To enable markerless insertion, the incorporation of self-excision cassettes commonly used in CRISPR/Cas9 and Flp/FRT systems may be useful. Last, the landing pad is marked with a *myo-2p::GFP* which is barely detectible under a dissecting microscope. Consequently, scoring for its disappearance after RMCE is occasionally difficult. This could be solved with a brighter fluorescent marker or stronger promoter.

### Dual-fluorescent operon reporter system

The ability to generate larger insertions (>10 kb) permits more complex transgenic experiments. As a proof of concept, we used RMCE to integrate dual fluorescence operon reporters for post-transcriptional gene regulation. Our experiments confirmed *let-7* regulation of the *lin-41* 3′-UTR in the seam cells and also revealed other potential regulation of the *lin-41* 3′-UTR (or possibly of the *unc-54* 3′-UTR) in other tissues.

Quantitative analysis of fluorescence protein amounts has long been employed to monitor gene expression and regulation in *C. elegans* (Breimann 2019). While initial approaches were largely limited to detecting the presence or absence of expression by location or time (Merritt *et al.* 2008), modern analysis techniques have become more sophisticated. Short half-life reporters (e.g., ~2 h for GFP/PEST) has allowed better quantification of temporal GFP expression in living worms (Frand *et al.* 2005). Another improvement has come with the advent of single copy insertion (e.g., using MosSCI (Frokjaer-jensen *et al.* 2008) and CRISPR/Cas9 (Dickinson and goldstein 2016)). For example, a quantitative two-color fluorescent reporter system was developed to more accurately measure in vivo images with one reporter containing a test 3′-UTR and the other containing the *unc-54* 3′-UTR that serves as the calibration control (Ecsedi *et al.* 2015). Although powerful, this design required placing two reporters driven by the same promoter, both in single copy, into two MoscSCI landing sites on different chromosomes, which is somewhat difficult (because of the construct size) and labor intensive. The phiC31 approach, which allows RMCE with a larger DNA fragment, reduces the time and labor bottleneck while also inserting both reporters into the same genomic site, which would remove any position effects. Thus, single copy integration by the phiC31 system should be of high value because it accelerates the examination of gene regulation, especially for long constructs such as dual fluorescence operon reporters.

### BAC integration challenge

We were unsuccessful with the clean integration of BAC sized constructs. Testing for recombination junctions in several strains revealed that many junctions were consistent with phiC31 mediated recombination, and these strains had lost phiC31. These results suggest that RMCE of an undesired insert occurred. The integrated DNAs were likely complex, possibly a product of the formation of jumbled extrachromosomal transgenes composed of rearrangements within the BAC and/or recombination with the co-injection marker prior to integration (Stinchcomb *et al.* 1985; Nonet 2020). Nevertheless, phiC31 mediated integration of BAC sized insertions may still be possible with additional optimization.

### Inversions

In addition to RMCE, we also used phiC31 integrase to create a large inversion on LG IV. The ability to generate precise inversions is useful in genetics and synthetic biology. In genetic analysis, large inversions can be used as genetic balancers, such as for maintaining strains carrying lethal mutations. Similarly, genetic balancers could be used to keep certain allele combinations together, as occurs naturally for selfish gene complexes, e.g., the *pha-1/sup35* maternal selfish element (Ben-david *et al.* 2017). Recently, a CRISPR/Cas9 approach was used to produce a powerful set of genetic balancers that spans about 89% of the *C. elegans* genome (Iwata *et al.* 2016; Dejima *et al.* 2018). By suitable placement of attP and attB sites, the phiC31 integrase system could be an alternative or additional approach for obtaining balancer for the remaining regions or for generating balancers over sub-regions.

In synthetic biology, recombinases have been used to control inversion switches in state machines or genome recorders (Friedland *et al.* 2009; Roquet *et al.* 2016). The detection of events in *C. elegans* has been limited to event reporting, such as by the induction of a fluorescent marker gene expression upon exposure to stresses (Link *et al.* 1999; Anbalagan *et al.* 2012). This approach is useful in situations when the signal, e.g., heavy metal presence, is occurring at the same time as, or possibly just prior to, observation. However, transient events that happened too far in the past cannot be recorded. Inversion recorders would fix the information into the genome such that prior exposure would be decoded later, even in subsequent generations. It would be interesting to utilize serine recombinases to record events in an animal, as has been done in microbial systems.

In summary, we have established phiC31 integrase RMCE as another option for precise single copy insertion of transgenes in *C. elegans.* Integrations are relatively easy to obtain for insert sizes up to ~15 kb. Although, our system is currently limited to one landing site, this site is suitable for quickly testing many constructs in the same genomic context and for inserting more complex genetic constructs. As an example, we have used the greater capacity to develop a dual operon reporter system to study post-transcriptional gene regulation. Finally, as expected, we have shown that the phiC31 integrase can be used to invert DNA sequences in *C. elegans.* Thus, the phiC31 integrase system should be a useful tool for genetic engineering, genetic analysis, and synthetic biology.

## Acknowledgements

We thank Dr. Helge Grosshans, Dr. Hitoshi Sawa, Dr. Chun-Liang Pan, Dr. Yi-Chun Wu and the *Caenorhabditis* Genetics Center (CGC) for providing *C. elegans* strains and plasmids. This work was supported by grants from the Ministry of Science and Technology, Taiwan (MOST 108-2311-B-002-012-), National Taiwan University Hospital (UN108-040) and National Taiwan University (109L7224) to SPC; and MOST (100-2311-B-001-015-MY3, MOST 104-2314-B-001-009-MY5) and Academia Sinica (103-CDA-L01, AS-GC-109-09) to JW.

